## Supplemental Figures for Figure 1-7 for "Defining cell type-specific immune responses in a mouse model of allergic contact dermatitis by single-cell transcriptomics"

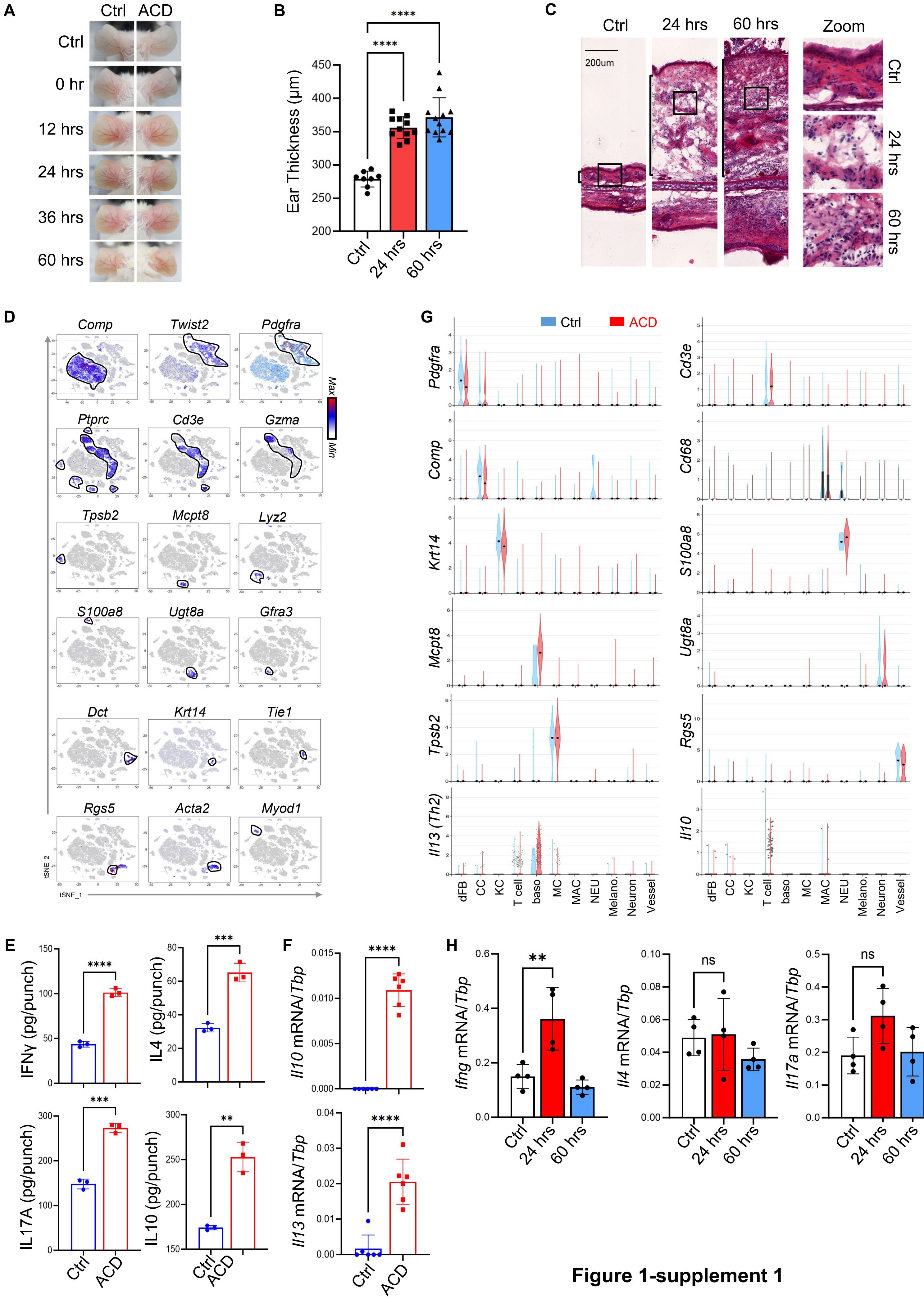


**Figure 1-supplement 1**

**Figure 1 - figure supplement 1: Establishment of the DNFB-elicited ACD mouse model**.
(**A**) Phenotypical presentation of mouse back skin after 0h, 12h, 24h, 36h and 60h of treatment.
(**B-C**) Ear skin samples from control, ACD (24 hours) and ACD (60 hours) groups were subjected for measurement of ear thickness (**B**) and histological HE analysis (**C**).
(**D**) tSNE plots showing the expression of indicated marker genes. Cell clusters expressing indicated markers were circled by black lines.
(**E**) Cell extracts isolated from control and ACD (60 hours) ear skin samples were subjected to ELISA analysis of IFNγ, IL4, IL17A and IL10 protein expression as indicated (n=3/group).
(**F**) Ear skin samples from control and ACD (60 hours) treatments were subjected to qRT-PCR analysis of *IL13, and Il10* mRNA expression as indicated (n=4~6/group).
(**G**). Violin plots showing the expression of indicated marker genes across various cell types, including *Pdgfra*$^{+}$*Comp*$^{-}$ dermal fibroblasts (dFB), *Comp*$^{+}$ chondrocytes (CC), *Krt14*$^{+}$ keratinocytes (KC), *Cd3*$^{+}$ or *Gzma*$^{+}$ T cells, *Mcpt8*$^{+}$ basophils (baso), *Tpsb2*$^{+}$ mast cells (MC), *Cd68*$^{+}$*Lyz2*$^{+}$ macrophages (MAC), *Ly6g*$^{+}$ neutrophils (NEU), *Dct*$^{+}$ melanocytes (Melano.), neuron-related cells (*Ug8a*$^{+}$ schwann cell and *Gfra3*$^{+}$ neuron), vessel-related cells (*Rgs5*$^{+}$ pericytes, *Acta2*$^{+}$ *Rgs5*$^{-}$ VSMC, and *Pecam1*$^{+}$ *Tie1*$^{+}$ endothelial cells) in ctrl and ACD samples.
(**H**). Skin draining lymph node samples from control and ACD (24 and 60 hours) –treated mice were subjected to qRT-PCR analysis of *Ifng*, *IL4, and Il17a* mRNA expression as indicated (n=4/group).

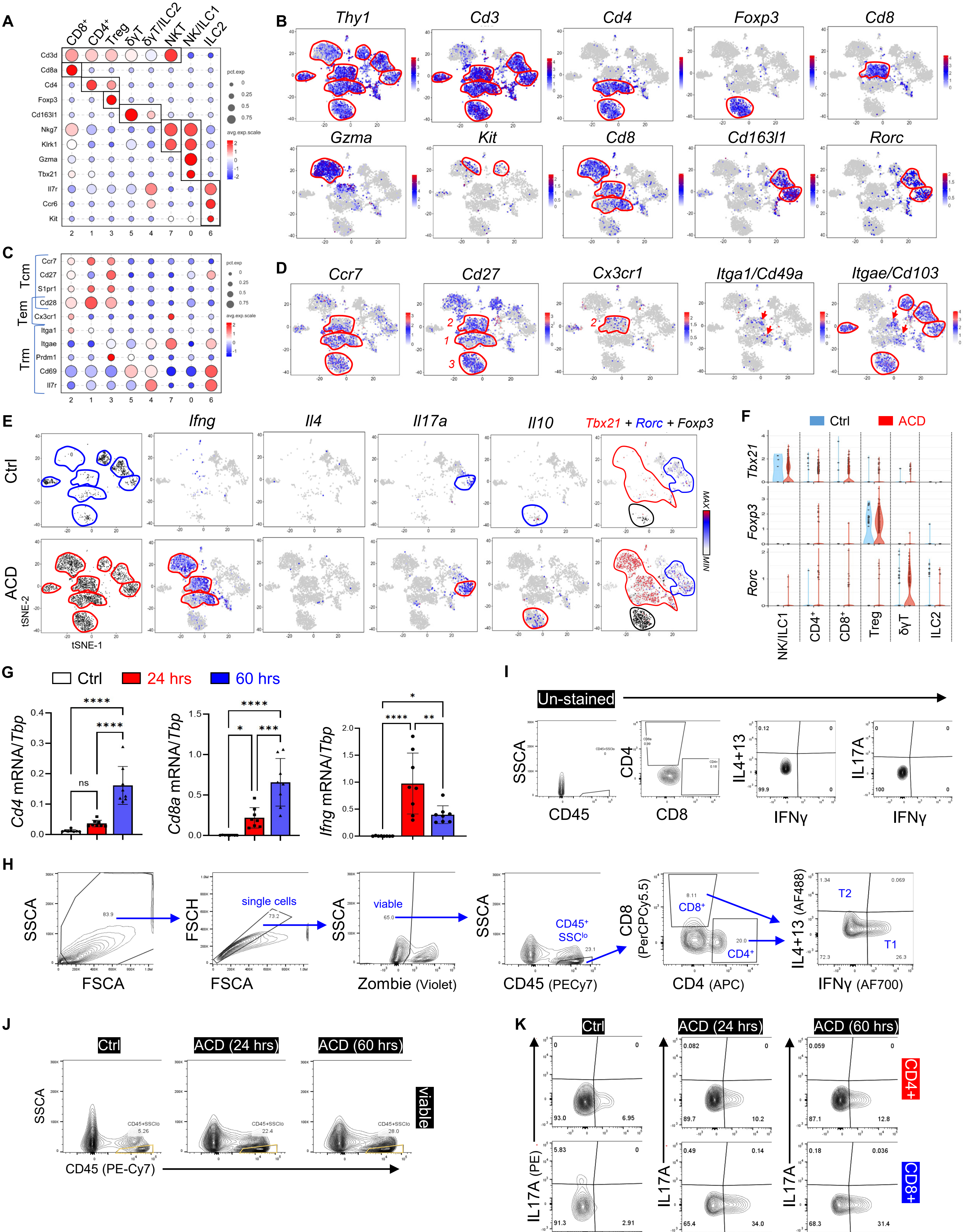


Figure 2-supplement 1

**Figure 2 - figure supplement 1: Characterization of T cell activation in ACD**.

(**A**) Bubble plots showing the expression of marker genes for each cell cluster. Abbreviations: *Treg*, regulatory T cell; *δγT*, delta gamma T cell; *ILC*, innate lymphoid cell; *NK*, natural killer cell; *NKT*, natural killer T cell.

(**B**) tSNE plots showing the expression of marker genes for each T cell cluster.

(**C-D**) Bubble plots (**C**) or tSNE plots (**D**) showing the expression of indicated marker genes associated with central memory (Tcm), effector memory (Tem) or tissue-resident memory (Trm) T cells for each cell cluster.

(**E**) tSNE plots showing cell distribution or the expression of *Ifng, Il4, Il17a,* and *Il10* in the control and the ACD samples. Far right is the overlaid tSNE plots showing the expression of *Tbx21* (red), *Rorc* (blue) and *Foxp3* (black) in T cell clusters in the control and the ACD samples.

(**F**) By sample violin plots showing the expression of indicated genes across various T cell sub-clusters.

(**G**) qRT-PCR analysis of the expression of *Cd4, Cd8a,* and *Ifng* in control, 24 hours, or 60 hours after elicitation of ACD (n=7~8/group). All error bars indicate mean ± SEM. *$p < 0.05$, **$p < 0.01$, ***$p < 0.001$, ****$p < 0.0001$, ns, non-significant.

(**H**) Gating strategies to analyze type 1 (T1) or type 2 (T2) $CD4^+$ or $CD8^+$ T cells in the skin.

(**I**) Unstained FACS plots for Fig. 2g.

(**J**) FACS plots showing the percentage of $CD4^+$ or $CD8^+$ T cells in $CD45^+SSCA^{lo}$ immune cells.

(**K**) FACS plots of IFNγ and IL17A in $CD4^+$ or $CD8^+$ T cells in control, ACD (24 hours) and ACD (60 hours) ear skin samples (n=3/group).

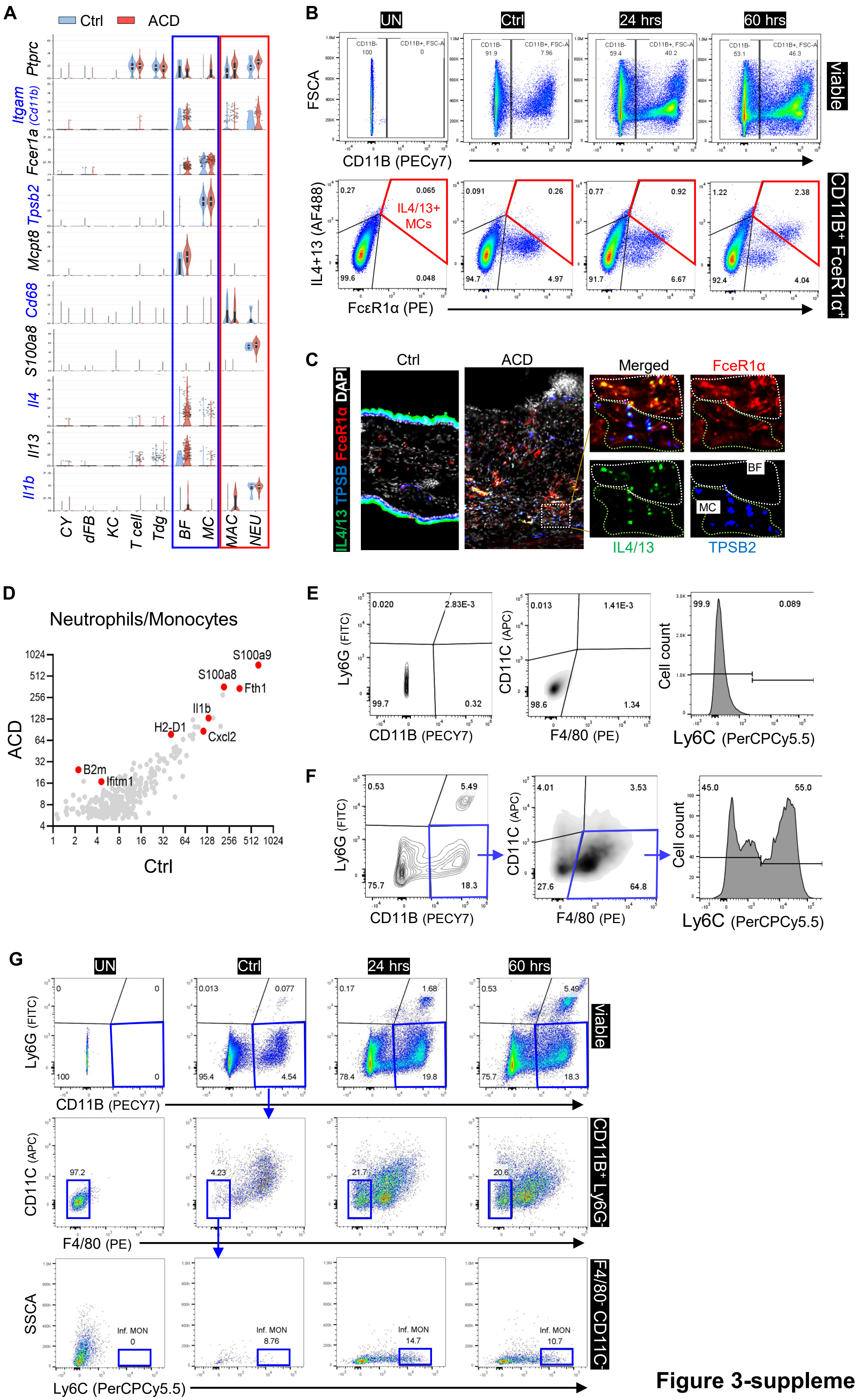


**Figure 3-supplement 1**

**Figure 3 - figure supplement 1: Analysis of myeloid cell activation in ACD**.

(**A**) Violin plots showing the expression of indicated genes across major skin resident and immune cells as indicated.

(**B**) FACS plots showing the percentage of $CD11B^-Fc\varepsilon R1a^+IL4/13^+$ (mast cell) or $CD11B^+$ $Fc\varepsilon R1a^+$ $IL4/13^+$ (basophil) cells within $CD45^+$ immune cells in control, ACD (24 hours) and ACD (60 hours) ear skin samples. Unstained (UN) plot was shown as negative gating control. All error bars indicate mean ± SEM. ****p < 0.0001.

(**C**) Frozen sections of control and ACD (60 hours) ear skin samples were subjected to immunostaining analysis using antibodies against IL4+IL13 (green), TBSP2 (blue) and FceR1a (red). Nuclei were counter stained by DAPI (white). Scale bar, 200 μm. Zoom-in image with overlaid or single-color channel is shown on the right hand side**.**

(**D**) Gene expression plot showing differentially expressed genes in control and ACD samples within the neutrophil cluster.

(**E**) Gating strategies to analyze Ly6C expression in $CD11B^+$;$F4/80^+$;$Ly6G^-$;$CD11C^-$ macrophages.

(**F**) unstained control plots for various markers used to analyze neutrophils or macrophages

(**G**) FACS plots showing the gating strategies to analyze $Ly6C^{hi}$ ($CD11B^+Ly6G^-F4/80^-CD11C^-$) inflammatory monocytes in viable cells

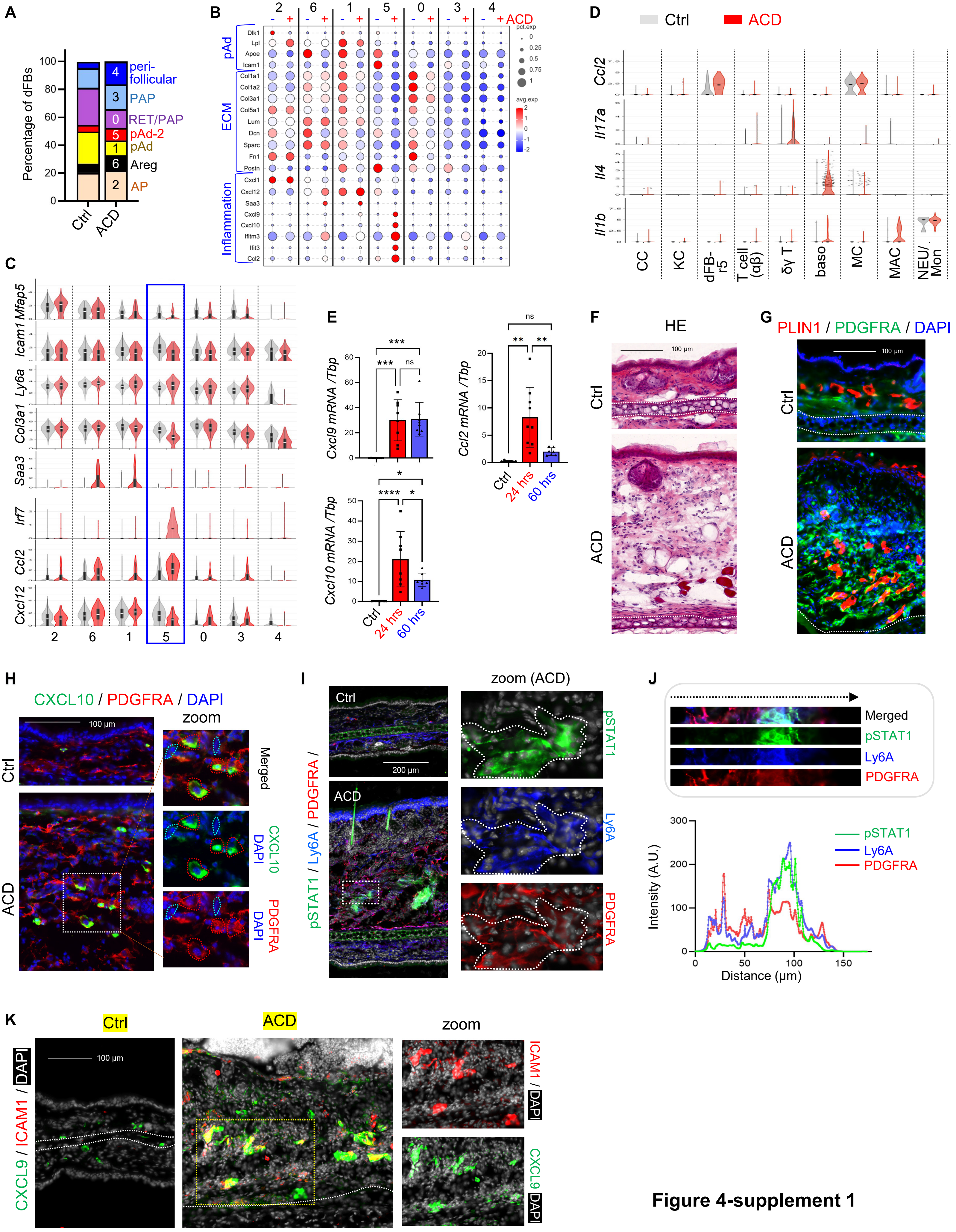


**Figure 4-supplement 1**

**Figure 4 - figure supplement 1: Characterization of the immune response of dermal fibroblasts in ACD**

(**A**) Stacked bar graphs showing the percentage of each dFB sub-cluster in the control and the ACD samples.

(**B-C**) Bubble plots (**B**) or Violin plots (**C**) showing the expression of indicated genes across various dFB sub-populations in the control and the ACD samples.

(**D**) Violin plots showing the expression of indicated genes across various cell populations in the control and the ACD samples.

(**E**) qRT-PCR analysis of the expression of *Cxcl9, Cxcl10*, *Ccl2* in control, 24 hours, or 60 hours after elicitation of ACD (n=7~8/group). Error bars indicate mean ± SEM. n.s., non-significant; **p < 0.01.

(**F**) Frozen sections of control and ACD ear skin samples were subjected to hematoxylin and eosin (HE) staining. Dash line marks the boarder between the dermis and the cartilage.

(**G**) Frozen sections of control and ACD ear skin samples were subjected to immunostaining analysis using antibodies against PDGFRA (green) and adipocyte marker PLIN1 (red). Nuclei were counter stained by DAPI (blue). Scale bar, 100 μm.

(**H**) Frozen sections of control and ACD ear skin samples were subjected to immunostaining analysis using antibodies against CXCL10 (green) and PDGFRA (red). Nuclei were counter stained by DAPI (blue). Scale bar, 100 μm. Zoom-in image is shown on the right-hand side, in which dermal $CXCL10^{+}$ or $PDGFRA^{-}$ cells were highlighted by either red- or blue-dotted lines.

(**I**) Frozen sections of control and ACD ear skin samples were subjected to immunostaining analysis using antibodies against pSTAT1 (green), Ly6A (blue) and PDGFRA (red). Nuclei were counter stained by DAPI (white). Scale bar, 200 μm. Zoom-in image is shown on the right-hand side.

(**J**) Quantified result showing the fluorescent intensity (arbitrary unit, AU) of pSTAT1 (green), Ly6A (blue) and PDGFRA (red) across image shown on the top panel from left to right.

(**K**) Frozen sections of control and ACD ear skin samples were subjected to immunostaining analysis using antibodies against CXCL9 (green) and ICAM1 (red). Nuclei were counter stained by DAPI (white). Scale bar, 100 μm. Zoom-in image of ACD sample is shown on the lower panel.

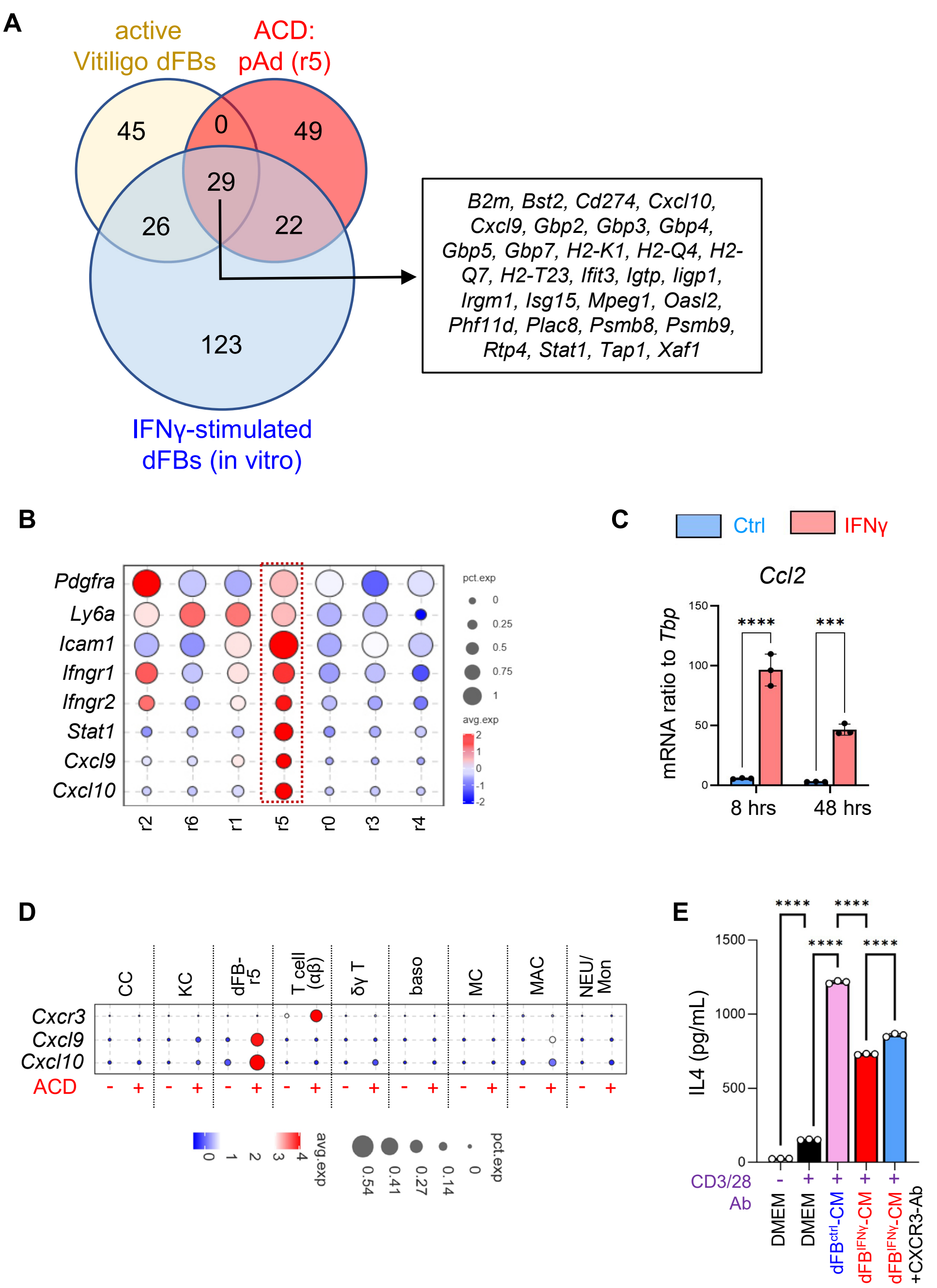


**Figure 5 - figure supplement 1: Interaction between dFBs and T cells via the IFNG-CXCL10-CXCR3 signaling axis in ACD.**

(**A**) Venn diagram comparing the top 100 IFNγ signaling dependent genes upregulated in the active vitiligo mouse dFBs (Xu et al. Nature 2022), the top 100 genes enriched in the pAd (dFB_r5) cells, and the top 200 IFNγ-inducible genes in primary dFBs (in vitro). The identities of the 29 genes upregulated in all 3 conditions are shown in the box on the right panel as indicated.

(**B**) Violin plots showing the expression of indicated genes across various dFB clusters

(**C**) Primary dFBs were treated with IFNγ for 8h or 48h, and control or IFNγ treated samples were subjected to qRT-PCR analysis of *Ccl2* mRNA expression (n=~3/group).

(**D**) Bubble plots showing the expression of indicated genes across various cell populations

(**E**). Primary naïve T lymphocytes were co-stimulated with CD3/CD28 antibody and control or IFNγ-primed dFB-CM with or w/o CXCR3 neutralizing antibody, and cell supernatants were collected for ELISA analysis of IL4 protein levels.

All error bars indicate mean ± SEM. **p < 0.01; ***p < 0.001; ****p < 0.0001; ns, non-significant.

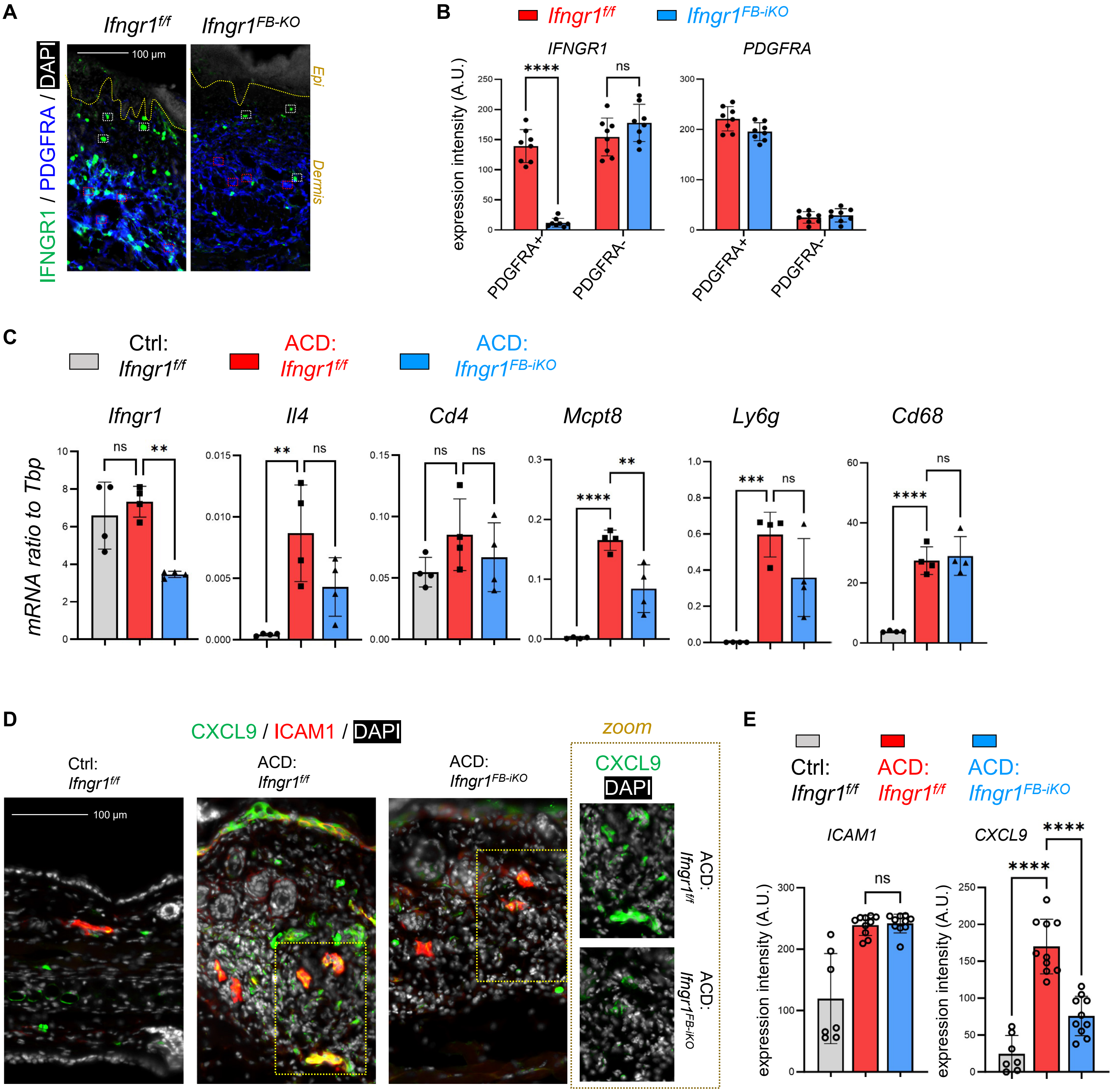


**Figure 6 - figure supplement 1: Targeted deletion of *Ifngr1* in dFBs inhibited the development of type-1 skin inflammation in ACD.**

(**A-B**) Lesional skin samples were immunostained with IFNGR1 and PDGFRA (**A**), and quantified result (**B**) showing the expression of IFNGR1 or PDGFRA in PDGFRA$^+$ dFB or PDGFRA$^-$ dermal cells as indicated. Yellow dash line indicated the epidermal-dermal junction; white boxes mark PDGFRA$^-$ dermal cells and red boxes mark PDGFRA$^+$ cells for quantification analysis (C); scale, 100 μm. All error bars indicate mean ± SEM. ****p < 0.0001; ns, non-significant.

(**C**) qRT-PCR analysis of the mRNA expression levels of indicated genes (ratios to HK gene *Tbp* were shown, n=3~4/group). All error bars indicate mean ± SEM. **p < 0.01; ***p < 0.001; ****p < 0.0001; ns, non-significant.

(**D-E**) Frozen sections of control and ACD ear skin samples were subjected to immunostaining analysis (**D**) using antibodies against CXCL9 (green) and ICAM1 (red). Nuclei were counter stained by DAPI (white). Scale bar, 100 μm. Zoom-in images are shown on the lower panel. Quantified result (**E**) showing the expression of ICAM1 or CXCL9 in ICAM1$^+$ cells as indicated.

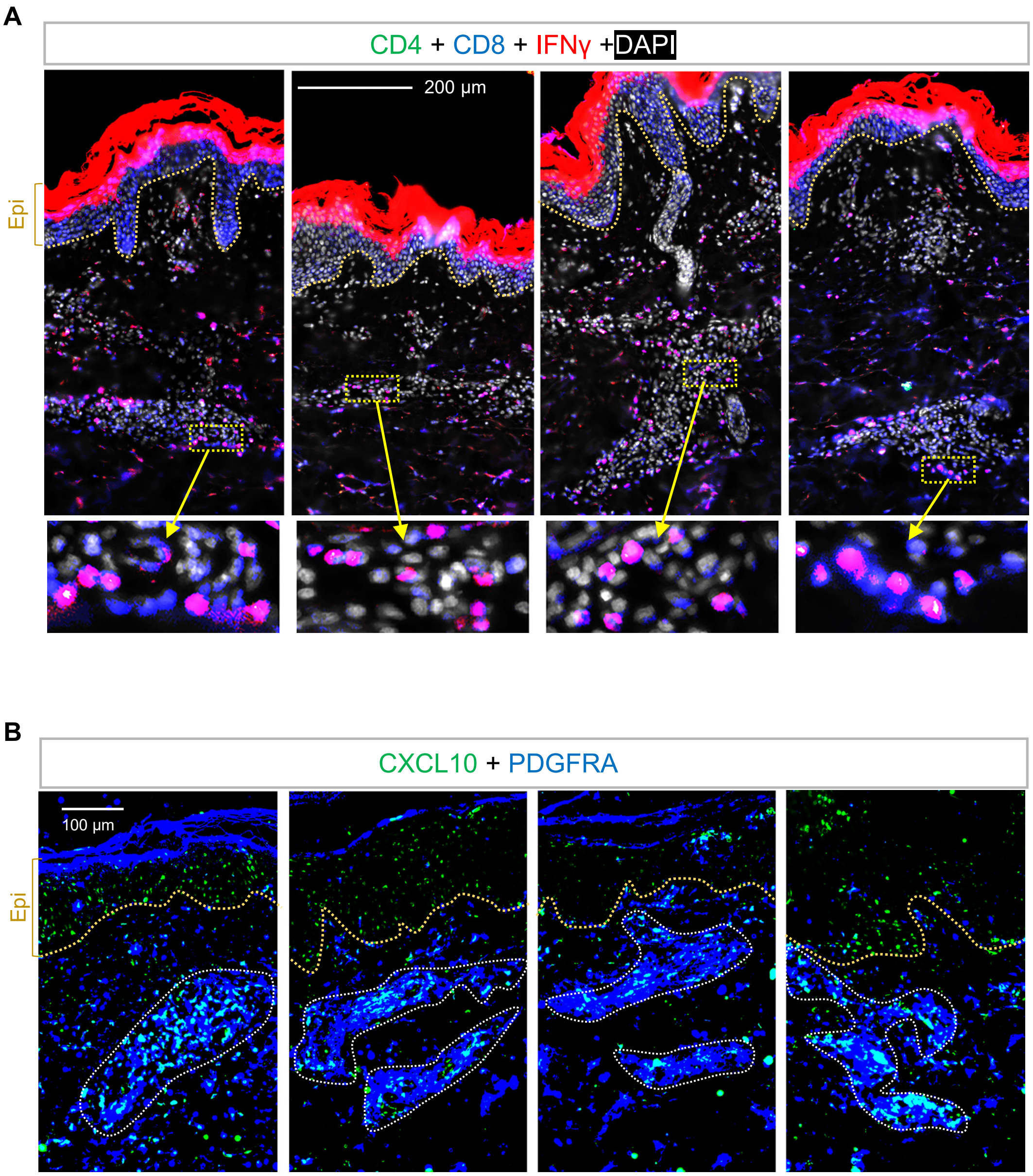


**Figure 7 - figure supplement 1: Activation of dermal T cells and fibroblasts in human ACD skin samples**.

(**A**) Skin sections from ACD human skin samples were immunostained with antibodies against CD4 (green), CD8 (blue) and IFNγ (red) (n=4). Nuclei were counter stained by DAPI (white). Scale bar, 200 μm. Zoom-in image is shown on the bottom panel.

(**B**) Skin sections from ACD human skin samples were immunostained with CXCL10 (green) and PDGFRA (blue) (n=4). Scale bar, 200 μm.
