## Supplementary Table S1 for "Defining cell type-specific immune responses in a mouse model of allergic contact dermatitis by single-cell transcriptomics"

**Table S1. List of gene primers used for RT-qPCR:**

| Gene | Strand | Primer sequence |
|---|---|---|
| *Tbp* | Forward | CCTTGTACCCTTCACCAATGAC |
| | Reverse | ACAGCCAAGATTCACGGTAGA |
| *Col1a1* | Forward | GCTCCTCTTAGGGGCCACT |
| | Reverse | ATTGGGGACCCTTAGGCCAT |
| *Il1b* | Forward | GAAATGCCACCTTTTGACAGTG |
| | Reverse | TGGATGCTCTCATCAGGACAG |
| *Ly6g* | Forward | GACTTCCTGCAACACAACTACC |
| | Reverse | ACAGCATTACCAGTGATCTCAGT |
| *Ifng* | Forward | GCCACGGCACAGTCATTGA |
| | Reverse | TGCTGATGGCCTGATTGTCTT |
| *Il4* | Forward | GAGCCATATCCACGGATGCGAC |
| | Reverse | ATGCGAAGCACCTTGGAAGCCC |
| *Il17a* | Forward | ACGCGCAAACATGAGTCCAGGG |
| | Reverse | TGAGGGATGATCGCTGCTGCCT |
| *Mcpt8* | Forward | AACGCTGAAGGAGGGGAAATCA |
| | Reverse | TTGCCACCAGGAAACCACCA |
| *Cma1* | Forward | CACGGAGTGCATACCACACT |
| | Reverse | AAGCTTCTGCCACGTGTCTT |
| *Cd68* | Forward | CTTCCCACAGGCAGCACAG |
| | Reverse | AATGATGAGAGGCAGCAAGAGG |
| *Cxcl9* | Forward | GGAGTTCGAGGAACCCTAGTG |
| | Reverse | GGGATTTGTAGTGGATCGTGC |
| *Cxcl10* | Forward | CCACGTGTTGAGATCATTGCCACG |
| | Reverse | ATCCATCGCAGCACCGGGGT |
| *Cxcr3* | Forward | TACCTTGAGGTTAGTGAACGTCA |
| | Reverse | CGCTCTCGTTTTCCCCATAATC |
| *Cd8a* | Forward | GACCGGATTGGACTTCGCCT |
| | Reverse | GACTAGCGGCCTGGGACATT |
| *Il10* | Forward | GGCGCTGTCATCGATTTCTCCCC |
| | Reverse | GGCCTTGTAGACACCTTGGTCTTGG |
| *Il13* | Forward | TGCTTGCCTTGGTGGTCTCGC |
| | Reverse | GCGGCCAGGTCCACACTCCA |
| *Cd4* | Forward | CTAGCTGTCACTCAAGGGAAGA |
| | Reverse | CGAAGGCGAACCTCCTCTAA |
| *Ccl2* | Forward | CACAGTTGCCGGCTGGAGCA |
| | Reverse | CAGCAGGTGAGTGGGGCGTT |
| *Stat1* | Forward | TCACAGTGGTTCGAGCTTCAG |
| | Reverse | CGAGACATCATAGGCAGCGTG |

| | | |
|---|---|---|
| ***Apoe*** | Forward | CTGACAGGATGCCTAGCCG |
| | Reverse | CGCAGGTAATCCCAGAAGC |
| ***Ifngr1*** | Forward | TGACTATGCACGGTCAAAAGAG |
| | Reverse | ATTCACAACGACTTCAGGGTG |
